# Sense the Moment: a highly sensitive antimicrobial activity predictor based on hydrophobic moment

**DOI:** 10.1101/2020.07.15.205419

**Authors:** William F. Porto, Karla C. V. Ferreira, Suzana M. Ribeiro, Octavio L. Franco

**Affiliations:** Porto Reports, Brasília-DF, Brazil; Pós-Graduação em Ciências Genômicas e Biotecnologia Universidade Católica de Brasília, Brasília-DF, Brazil; Centro de Análises Proteômicas e Bioquímicas, Pós-Graduação em Ciências Genômicas e Biotecnologia Universidade Católica de Brasília, Brasília-DF, Brazil; Programa de Pós-Graduação em Ciências da Saúde, Universidade Federal da Grande Dourados, Dourados, MS, Brazil; S-Inova Biotech, Pós-Graduação em Biotecnologia, Universidade Católica Dom Bosco, Campo Grande, MS, Brazil

**Keywords:** Shuffled Peptides, Antimicrobial Peptides, Hydrophobic Moment, Peptide screening, α-Helical Peptides

## Abstract

**Background:** Computer-aided identification and design tools are indispensable for developing antimicrobial agents for controlling antibiotic-resistant bacteria. Antimicrobial peptides (AMPs) have aroused intense interest, since they have a broad spectrum of activity, and therefore, several systems for predicting antimicrobial peptides have been developed, using scalar physicochemical properties; however, regardless of the machine learning algorithm, these systems often fail in discriminating AMPs from their shuffled versions, leading to the need for new training methods to overcome this bias. Aiming to solve this bias, here we present “Sense the Moment”, a prediction system capable of discriminating AMPs and shuffled versions.

**Methods:** The system was trained using 776 entries: 388 from known AMPs and another 388 based on shuffled versions of known AMPs. Each entry contained the geometric average of three hydrophobic moments measured with different scales.

**Results:** The model showed good accuracy (>80 %) and excellent sensitivity (>90 %) for AMP prediction, exceeding deep-learning-based methods.

**Conclusion:** Our results demonstrate the system’s applicability, aiding in identifying and discarding non-AMPs, since the number of false negatives is lower than false positives. General Significance: The application of this model in virtual screening protocols for identifying and/or creating antimicrobial agents could aid in the identification of potential drugs to control pathogenic microorganisms and in solving the antibiotic resistance crisis.

**Availability:** The system was implemented as a web application, available at <http://portoreports.com/stm/>.

## 1 Introduction

In the last few decades, the antibiotic resistance crisis has led to the need for new antimicrobial compounds because the activity of conventional antibiotics has been reduced. In this context, antimicrobial peptides (AMPs) have been proposed as an alternative strategy to control antibiotic-resistant microorganisms.^1–3^ AMPs are evolutionarily ancient molecules that have been identified from diverse sources, such as microorganisms, plants and animals.^4–7^ They play an important role in the innate immune system and are the first line of defense to protect internal and external surfaces of the host.^7^ Overall, AMPs can adopt amphipathic structures, with a positive net charge, and they are composed of dozens of amino acid residues.^8^ AMPs comprise a highly diverse group of antimicrobial compounds, split into two major groups, according to the presence or absence of disulphide bonds: cysteine-stabilized and disulphide-free peptides, respectively.^8^ In addition, AMPs can be classified according to their origins: naturally occurring AMPs, which can be found in members of all kingdoms of life, and which are a product of specific genes;^4–7^ cryptic AMPs, in larger proteins, where proteolytic cleavage under specific environmental conditions releases them;^9–11^ and designed AMPs, developed with recent technologies of rational design and peptide synthesis.^1,12,13^

Conducting a search for AMPs in databases and undertaking their rational design remains a promising method for identifying and/or creating novel antimicrobial agents against pathogenic microorganisms. However, the identification of an active peptide sequence prior to chemical synthesis and susceptibility testing represents a challenge. Indeed, the identification prior to synthesis requires sophisticated techniques, since tens^12^ to hundreds^13^ of peptides still need to be synthesized and assayed for their antimicrobial potency. Therefore, prediction systems could help to reduce the number of peptides to be tested, especially those systems implemented following Chou’s 5-step rule.^14^ Most AMP prediction systems have the same or similar prediction assessments,^17^ regardless of machine learning algorithm (e.g. neural network, support vector machine), because they have been based on physicochemical/structural properties, such as charge, hydrophobicity and secondary structure formation.^8,15,16^

The application of physicochemical/structural properties introduces a bias to prediction, which was demonstrated by the linguistic model for the rational design of antimicrobial peptides.^12^ This model showed up a problem that hinders correct classification in AMP prediction systems: shuffled versions of AMP sequences have the same physicochemical properties (e.g. charge and hydrophobicity), but the antimicrobial activity is abolished.^12^ In this context, our group previously demonstrated that AMP prediction tools often fail when challenged with shuffled sequences, even in the case of iAMP-2L, which uses the correlation between amino acid residues based on Chou’s pseudo amino acid composition.^17^ Therefore, exploring genomes and/or sequence databases with current antimicrobial activity prediction tools is not feasible, because sequences with a similar composition could generate a huge number of false positive hits.

In this context, in 2010, our group proposed the inclusion of vector properties, such as hydrophobic moment, as developed by Eisenberg et al.,^18^ as a way to avoid this bias of identical composition of shuffled sequences.^19^ However, although the hydrophobic moment is strongly connected with the sequence, this property is not correlated with antimicrobial activity.^20,21^ Despite that, we hypothesized that the hydrophobic moment could be used as a classifier between ordered and shuffled sequences. Therefore, here we report the development of a sensitive system for distinguishing ordered and shuffled peptides, with high sensitivity for AMP prediction.

## 2 Results

### 2.1 Hydrophobicity scale influences the hydrophobic moment

Because the hydrophobic moment depends on the hydrophobic scale, we measured the pairwise root main squared error (RMSE) and the Euclidian distance between the calculated hydrophobic moments for each sequence in the positive dataset (Table 1). The Eisenberg and Wimley-White scales showed a similar trend, presenting the lowerest values for RMSE and Euclidian distance (Table 1). Therefore, for this study we selected the Eisenberg over the Wimley-White scale, because the Eisenberg scale is a consensus scale. Therefore, for the next steps, the scales from Eisenberg, Kyte-Doolittle and Radzicka-Wolfenden were used for hydrophobic moment calculation.

**Table 1.**
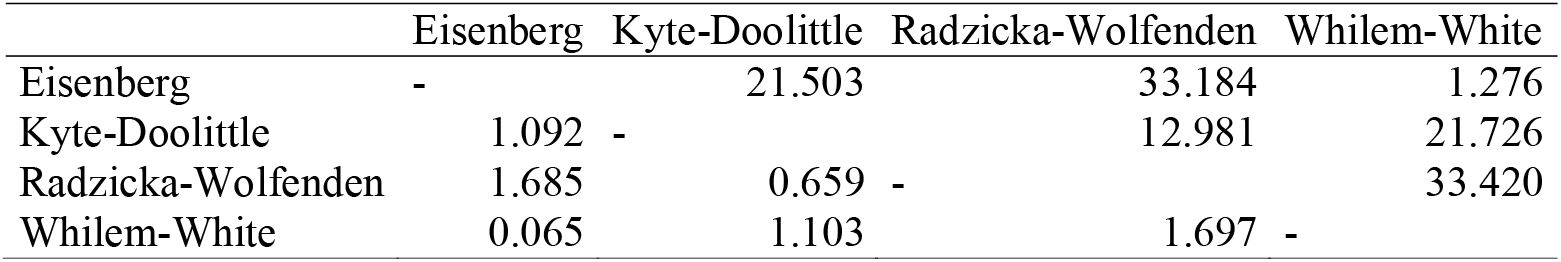
Pairwise distances between the hydrophobic moments calculated with different hydrophobicity scales. The upper triangle shows the Euclidian distance, while the lower triangule shows the RMSE values.

### 2.2 Hydrophobic moment discriminates AMPs and their shuffled versions

In order to discriminate shuffled sequences, the sequences described by Loose et al.^12^ would be used as training data, because this data set is the proof of principle that shuffled sequences have lost their activities. However, this data set has few sequences for training the system, and we preferred to use these sequences as a blind set for testing the system. Therefore, instead of using the Loose data set, we used APD sequences as the positive data set (PS); and as the negative data set (NS), we used a theoretical set, generated by shuffling sequences from the positive data set (Figure 1A). The datasets are available in supplementary Tables 1-3.

**Figure 1.**
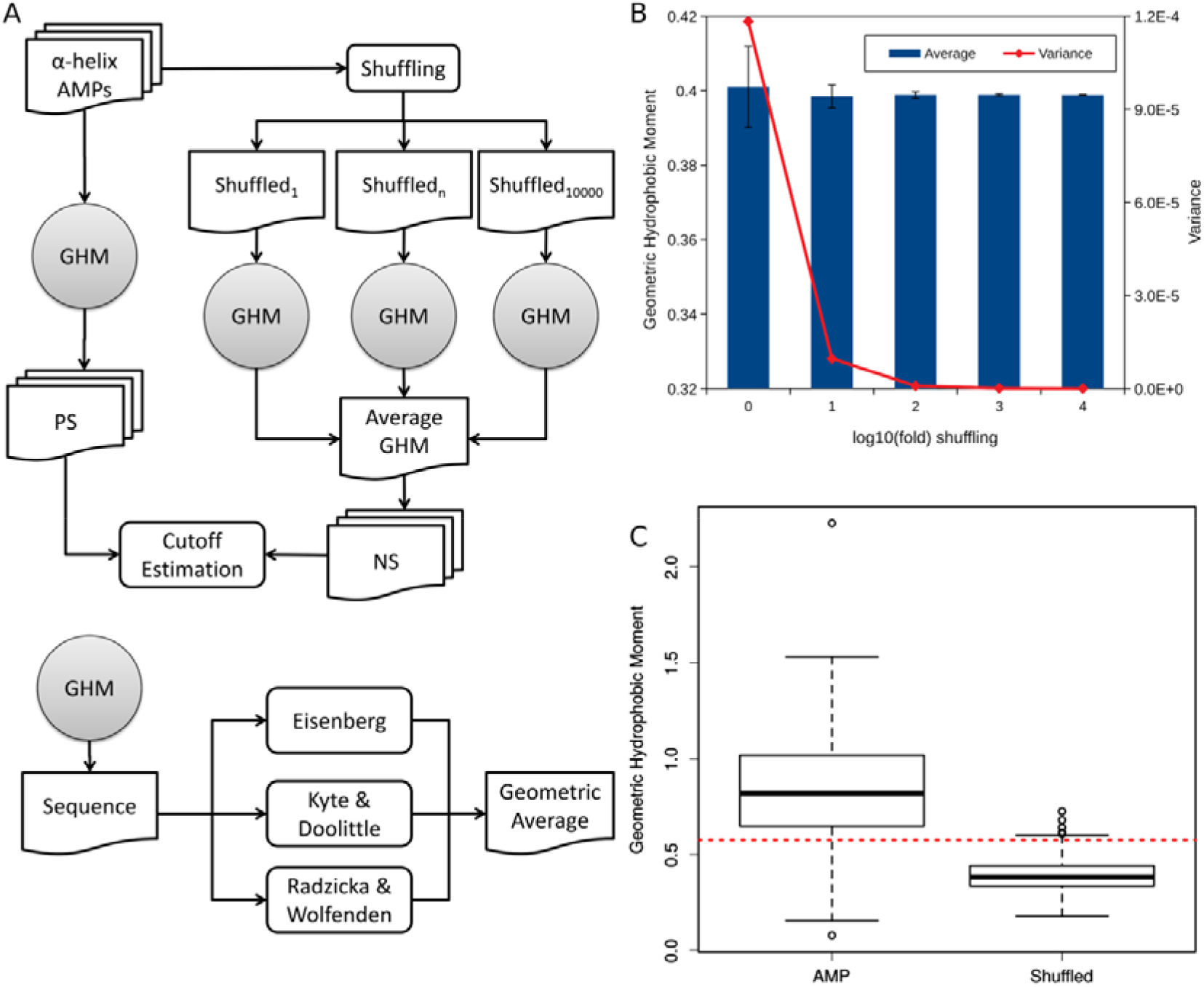
Discriminating AMPs and their shuffled versions using the geometric hydrophobic moment. (A) The dataset construction flowchart. The α-helical peptides were subjected to geometric hydrophobic moment calculation, generating the positive dataset (PS). The same set of peptides was subjected to a shuffling process in order to generate the negative dataset (NS). A total of 10,000 rounds of shuffling were applied in order to abolish the variance generated by the randomization process. Then, PS and NS were used for threshold definition. Because the geometric hydrophobic moment (GHM) is a single value, there is no need to use any machine learning engine for classification. (B) Reducing data variance from shuffled sequences. Data represent the average of 100 independent shuffling experiments. In each experiment, each sequence was shuffled n-folds, where n is expressed as the log_10_(fold) shuffling. Since our negative data set was composed of shuffled sequences, increasing the number of shuffling rounds reduces the variance in the set, allowing reproducibility, even with the bias of randomness data. Shuffling the peptides by 10,000 folds almost abolished the variance and consequently the standard deviation. (C) Boxplot of geometric hydrophobic moment distribution within PS and NS. Despite the overlap between the sets, the geometric hydrophobic moment values calculated from PS were statistically higher than NS according to the Wilcoxon-Mann-Whitney non-parametric test, showing the same p-value for the three used scales (p-value <2.2e-16). In fact, only 24 out of 388 shuffled sequences (6.19 %) had their geometric hydrophobic moments increased after shuffling. The dotted red line indicates the best cut-off value (0.590).

For PS generation, initially, all sequences with at least 70 % of helical content on PSIPRED predictions were selected, then the redundant sequences at 70 % were removed, and finally, the set was curated by hand, with 388 sequences remaining (Supplementary Table S1).

For NS generation, a single round of shuffling did not ensure the generation of a shuffled peptide, especially if the original sequence was composed of a set of restricted and repetitive amino acids. In addition, being a random process, shuffling increases the dataset variance, which could hinder reproducibility. Therefore, to reduce the variance and in order to obtain homogeneous hydrophobic moment values, the shuffling process was repeated 1, 10, 100, 1,000 and 10,000 times for each sequence, and then the set was averaged (Figure 1). Although there is no statistical difference between the sets regardless of the repetition times, increasing the repetitions abolished the variance caused by shuffling, and the set with 10,000 repetitions was selected.

Because we are assessing a single descriptor, there is no need for a machine learning classifier. Overall, geometric hydrophobic moment values in NS were reduced in comparison with PS (Figure 1C). A similar trend was observed using individual scales (data not shown). According to the receiver-operating characteristic (ROC) curve (Figure 2), the system presented a very good performance, showing an area under the curve of 0.891, where the threshold of 0.59 showed an accuracy of 88.79 %. The thresholds for single scales are 0.19, 0.83 and 1.28 for, respectively, Eisenberg, Kyte-Doolittle and Radzicka-Wolfenden scales. By using a single scale instead of the GHM, the results are similar; however, GHM acts as a voting system (i.e. the sequence KRFGRLAKSFLRMRILLPRRKILLAS was predicted as AMP by Radzicka-Wolfenden and Eisenberg scales but not by Kyte-Doolittle scale).

**Figure 2.**
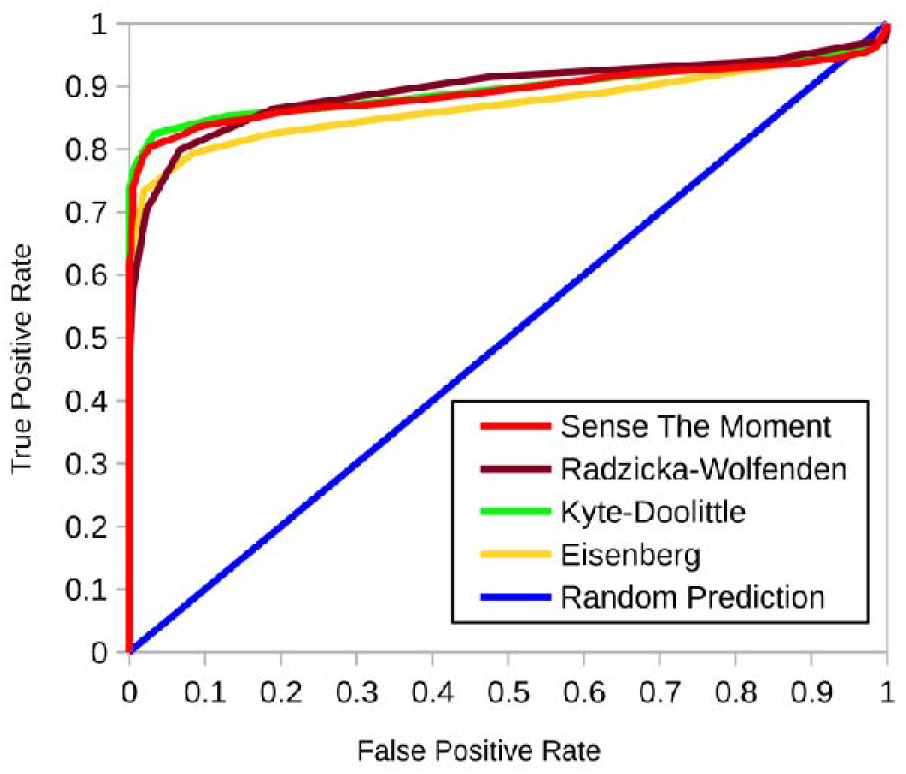
Receiver-operating characteristic (ROC) curve concerning the training set. Sense the Moment, Radzicka-Wolfenden, Kyte-Doolittle, Eisenberg and Random predictions are represented by, respectively, red, brown, green, yellow and blue lines. The AUC values were, respectively, 0.891, 0.895, 0.895, 0.869 for Sense the Moment Prediction, Radzicka-Wolfenden, Kyte-Doolittle, Eisenberg predictions. The results are consistent with the 5-fold cross-validation.

### 2.3 Sense the Moment is Comparable to Deep learning approaches

Tables 2 and 3 summarise the assessments of Sense the Moment against blind data sets from Loose and Nagarajan, respectively. The first evaluation, taking into account only the training data, slightly overestimated the model, because accuracies in Loose and Nagarajan data sets were 80.77 and 85 %, respectively. Notably, sensitivity drew attention, due to excellent rates in both blind sets, reaching 95 and 94.4 % in Loose and Nagarajan data sets, respectively. However, if the MCC value was taken into account, the system was better in discriminating designed and shuffled peptides than antimicrobial and non-antimicrobial peptides. Furthermore, the model has a PPV of 74.51 and 89.47 % in Loose and Nagarajan data sets, respectively. Interestingly, methods based on the hydrophobic moment, Sense the Moment and DBAASP, outperformed the systems based on deep learning algorithms AMP Scanner Vr. 2 and Deep-AmPEP. DBAASP showed the best performance against Loose and Nagarajan data sets, while Sense The Moment was ranked 2^nd^ in Loose (Table 2); and 3^rd^ in the Nagarajan data set (Table 3).

**Table 2.** Prediction assessments of Sense the Moment and its benchmarking against Deep learning methods in the Loose data set.

| Algorithm | Sensitivity | Specificity | Accuracy | PPV | MCC |
| --- | --- | --- | --- | --- | --- |
| Sense The Moment | 95.00 | 65.79 | 80.77 | 74.51 | 0.64 |
| DBAASP | 100.00 | 73.68 | 87.18 | 80.00 | 0.78 |
| Deep-AmPEP | 77.50 | 23.68 | 51.28 | 51.67 | 0.01 |
| AMP Scanner Vr. 2 | 100.00 | 36.84 | 69.23 | 62.60 | 0.48 |

**Table 3.** Prediction assessments of Sense the Moment and its benchmarking against Deep learning methods in the Nagarajan data set.

| Algorithm | Sensitivity | Specificity* | Accuracy | PPV | DOR** |
| --- | --- | --- | --- | --- | --- |
| Sense The Moment | 94.44 | 0.00 | 85.00 | 89.47 | 2.33 |
| DBAASP | 88.89 | 50.00 | 85.00 | 88.89 | 6.60 |
| Deep-AmPEP | 88.89 | 50.00 | 85.00 | 94.12 | 6.60 |
| AMP Scanner Vr. 2 | 83.33 | 0.00 | 75.00 | 88.24 | 0.89 |

\* There are only two negative entries in the set, thus, specificity would be 0, 50 or 100%.
\*\* Because of the high prevalence of positive entries in this data set (90 %), DOR was used as a prevalence independent measurement.

## 3 Discussion

In the last decade, numerous computational methods to predict the antimicrobial activity of AMPs have been reported.^8,15,16,19,21–23^ The majority of these methods use different machine learning algorithms; however, this results in only a slight variation in prediction performance. Besides, these prediction tools are not able to handle shuffled sequences.^17^ Because a shuffled sequence has the same scalar properties as the native sequence, the sequences would have the same or similar predictions.^17^ Thus, any arbitrary sequence with a similar composition to an AMP would be predicted as an AMP. Hence, considering the choice of a machine learning algorithm with little or no effect on prediction performance, training is the Achilles’ heel of all these systems. Most systems use scalar properties for training algorithms,^8,15,16^ resulting in the compositional bias.^17^ This is a problem when genomes and databases are used as a source for such molecules.^24^

Therefore, the use of descriptors that take the amino acid order into account is desirable to discriminate between AMPs and non-AMPs with similar sequences.^19^ In this context, there are some properties that change when sequence order is altered, including hydrophobic moment^18^ and Chou’s pseudo amino acid composition.^25^ The iAMP-2L server^26^ was constructed using Chou’s pseudo amino acid composition; however, in our previous report, this server was not able to distinguish between designed and shuffled sequences.^17^ Currently, deep learning is the state of the art in terms of algorithms for prediction, and this approach is capable of extracting data direct from sequences, with no need to convert the sequence into descriptors.^27,28^ However, the number of sequences required for training is a clear limitation for deep learning application to AMPs and, therefore, this could be the main reason why Deep-AmPEP and AMP Scanner Vr. 2 were outperformed by hydrophobic moment-based systems (Tables 2 and 3).

Here, we posited hydrophobic moment as a way to distinguish designed and shuffled sequences, because this property is almost a fingerprint of amino acid order. Thus, assuming α-helical AMPs have higher hydrophobic moment values, shuffled sequences would have their hydrophobic moments reduced. Indeed, it is only in 6.19 % of AMP sequences that the geometric hydrophobic moment increases when the sequence is shuffled and, therefore, there are statistical differences in hydrophobic moments, when comparing AMPs and shuffled peptides (Figure 1C).

However, there are two biases associated with the hydrophobic moment, since its value is connected, firstly, to the hydrophobicity scale used^29^ and, secondly, to the peptide’s secondary structure.^20^ Although hydrophobicity scales are strongly correlated,^29–33^ the hydrophobicity of a given amino acid could differ within scales. For instance, in Eisenberg’s scale, phenylalanine is more hydrophobic than leucine, in contrast to Kyte-Doolittle’s. The differences between the scales may be due to reasons such as pH or experimental system. This fact is observed when the Euclidian distance and the RMSE between hydrohphobic moments are calculated with different scales (Table 1). To overcome this bias, Sense the Moment was developed using the geometric average of three hydrophobic moments, each one calculated with a different scale; and the geometric average was essential because the three values have the same weight independently of the scale used. In practical terms, the individual scales have similar predictive power (Figure 2). However, the use of the geometric average is important for those sequences which are near the threshold value, because the small differences between the scales could influence the prediction of these sequences. Finally, due to the strong correlations between the scales, the choice of scales makes only a small contribution to the results.^31^ Therefore, we applied the three scales which presented great differences in the hydrophobic moments according to Euclidian distance and RMSE : Eisenberg,^18^ Kyte-Doolittle^34^ and Radzicka-Wolfenden^35,36^ (Table 1).

The second bias, related to secondary structure, was handled by selecting AMP sequences with an α-helix as secondary structure, because δ in eq. 1 was set to 100°. Even so, Sense the Moment could, in theory, be applied to β-stranded peptides. Mathematically, β-strand structure could present elevated values of hydrophobic moment even with δ adjusted to an α-helix, because in an amphipathic α-helix, residues X_i_ and X_i+4_ would be on the same side (e.g. X_1_ and X_5_), while in an amphipathic β-strand, the residues on the same side are X_i_ and X_i+2_. Following this reasoning, in a β-strand, residues X_i_, X_i+2_ and X_i+4_ would be on the same side and, therefore, generating elevated values for the β-strand even with δ adjusted to an α-helix. In fact, in the Loose and Nagaranjan data sets some sequences were unable to form α-helixes.^17,37^ Therefore, Sense the Moment is restricted to short peptides with low-complexity secondary structures, in particular disulphide-free peptides.

In fact, the system had very good accuracy in both blind sets, considering designed/shuffled (Table 2) and AMP/non-AMP (Table 3). However, the strength of the system relies on its sensitivity, which is what gives the system its name. Because Sense the Moment has excellent sensitivity and only reasonable specificity, a few false negative predictions were expected. With this in mind, Sense the Moment should be used for removing peptides predicted as non-AMP, reducing the number of sequences to evaluate *in vitro* or even used in combination with other prediction tools.

However, the main limitation of Sense the Moment is that it merely takes the geometric hydrophobic moment into account, which has only two dimensions, while the three-dimensional structure of AMPs could be more complex and might interfere in antimicrobial activity. In fact, this could be the reason why the DBAASP predictor outperformed Sense the Moment in both datasets (Tables 2 and 3). DBAASP takes into account charge density and depth-dependent potential in addition to the hydrophobic moment itself. Interestingly, the DBAASP predictor and Sense the Moment were constructed using a different database, which confirms the binary prediction capacity of hydrophobic moment and also validates our approach for the construction of a negative data set (Figure 1A and 1B). However, we have to remember that the hydrophobic moment works best in classifying shuffled and designed sequences, not AMPs themselves, because not all designed sequences from Loose were indeed active against bacteria.^12^ In addition, there is no direct correlation between hydrophobic moment and antimicrobial activity.^20,21^

Therefore, Sense the Moment could be used together with other prediction systems, such as AmPEP and AMP Scanner, allowing the best characteristics of each system to be exploited, in particular with the application of Sense the Moment RESTFull API, which opens the possibility of generating consensus methods for AMP prediction.

## 4 Conclusions

The application of computers for identifying and/or designing AMPs has helped the field to move forward. These computational models can explore vast quantities of data and generate the next generation of AMPs. However, this is a recent field in machine learning applications, with nebulous and unexplored regions. At the beginning, the main biases were sequence length variation and the absence of a non-AMP database. By now, the main bias is discriminating AMPs from sequences with the same composition in a different order. Due to the small overlap between AMPs and their shuffled versions, Sense the Moment partially solved this bias. Henceforth, Sense the Moment could be a valuable tool for identifying drug candidates to help to solve the antibiotic resistance crisis.

## 5 Material and Methods

### 5.1 Hydrophobicity Scale Selection

Based on the positive data set of antimicrobial peptides (see below), the hydrophobic moment was measured using four different hydrophobic scales: Eisenberg,^18^ Kyte-Doolittle,^34^ Radzicka-Wolfenden^35,36^ and Wimley & White^38^ (Figure 1A). The average hydrophobic moment was calculated using Eisenberg’s equation:^18^

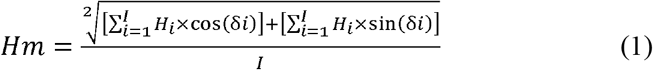

Where δ represents the angle between the amino acid side chains (100° for α-helix, on average); *i* represents the residue number in the position *i* from the sequence; H_*i*_ represents the *i*^th^ amino acid’s hydrophobicity on a hydrophobicity scale; and *I* represents the total number of residues.

Since the hydrophobic scales are correlated and although their values are numerically different, the resulting hydrophobic moment could present a similar trend; therefore, a pairwise comparison was performed by calculating the root mean squared error and the Euclidean distance between sets of hydrophobic moment values for each scale.

### 5.2 Data Sets

#### 5.2.1 Positive Data Set

The Antimicrobial Peptides Database^39^ was used as the primary source of AMP sequences. Initially, the secondary structure of all sequences was predicted using PSIPRED,^40^ and for the construction of the positive data set (PS), only sequences with at least 70 % of helical content were selected. Then, sequences with redundancy greater than 70 % were removed through CD-HIT.^41^ This set was manually curated, removing incomplete sequences, totalling 388 sequences. For each sequence, three average hydrophobic moments (equation 1) were calculated, using the Eisenberg,^18^ Kyte-Doolittle,^34^ and Radzicka-Wolfenden^35,36^ hydrophobicity scales. Then, the three hydrophobic moments were used to calculate the geometric average of the hydrophobic moment, which is referred to as the “geometric hydrophobic moment”.

#### 5.2.2 Negative Data Set

Given that shuffled versions of AMPs often lose their activity, the negative data set (NS) was derived from the positive data set by shuffling sequences (Figure 1A). Because shuffling is a random process, variance must be reduced to ensure reproducibility. Hence, an initial analysis was performed to verify the number of repetitions necessary to minimize this variance (Figure 1B). Ten thousand shuffled versions of each positive sequence were generated. The NS was composed of 388 entries, representing the average of 10,000 geometric hydrophobic moment values of shuffled versions of AMP sequences. The shuffling was performed using a PERL script, available at Git Hub (http://www.github.com/williamfp7/stm). A one-sided Wilcoxon-Mann-Whitney non-parametric test was applied to verify the differences between geometric hydrophobic moment in PS and NS sets, with a critical value of 0.05, using R package for statistical computing (http://www.r-project.org).

### 5.3 Prediction Cut-off Estimation

By using positive and negative sets together, the cut-off value was evaluated in intervals of 0.02, where for each interval true positive and true negative rates were measured. The cut-off value with the most correct predictions was selected as default. The area under the curve (AUC) was measured using the rollmean function of the zoo library from R package for statistical computing. For comparison, the same process was applied using the hydrophobic moments calculated using a single scale instead of the geometric hydrophobic moment.

#### 5.3.1 Blind Data Sets

Two blind data sets were used to assess the predictive capacity of Sense the Moment, regarding the differentiation between designed and shuffled sequences and the differentiation between antimicrobial and non-antimicrobial peptides. The first blind data set was derived from Loose et al.^12^ (Loose data set). The set was manually curated, removing sequences that had not been experimentally validated, resulting in 40 designed peptides and 38 shuffled peptides. The second blind data set was derived from Nagarajan et al.^37^ (Nagarajan data set), corresponding to 20 peptides tested against 30 organisms of Gram-negative, Gram-positive, mycobacterial, and fungal origin, where 18 were active against at least one organism and 2 were completely inactive.

### 5.4 Assessments against blind sets

The prevalence of both blind sets was measured according to Eq. 2:

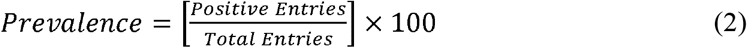

The prevalence indicates how often AMPs occur in the data set and could be used as an indication of how statistical tests should be used. The model was challenged against the blind data sets and the following parameters were measured:

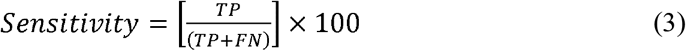

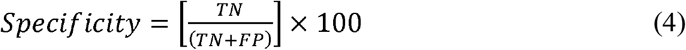

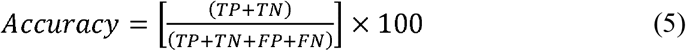

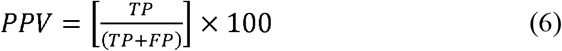

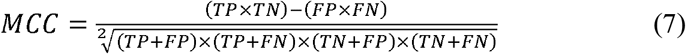

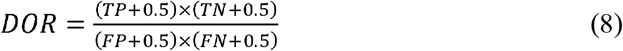

Where TP is the number of true positives; FN, the false negatives; TN, the true negatives; FP, the false positives; PPV, the probability of correct positive prediction; and MCC, Matthews Correlation Coefficient, which measures the quality of the prediction (the higher the value, the better the prediction is). Finally, the diagnostic odds ratio (DOR) is a prevalence independent indicator of performance, ranging from 0 to infinity, where values lower than 1 indicate an inverse prediction; values equal to 1, a random prediction; and values higher than 1, a correct prediction, with higher values indicating better discriminatory test performance.^42^ Since some divisions by zero may occur, the approximated DOR is calculated by adding 0.5 to all counts.^42^

### 5.5 Benchmarking

The performance of Sense the Moment was compared with the state-of-the-art methods for antimicrobial activity prediction, AMP Scanner Vr. 2^27^ and Deep-AmPEP^28^, which are based on deep learning, using Eqs. 3–7. In addition, the DBAASP prediction tool,^43^ which is based on the hydrophobic moment, charge density and depth-dependent potential, was also assessed against the blind data sets.

### 5.6 Web interface and RESTFull API

Sense the Moment was developed as a web application, using the PHP programming language. The system can be accessed by a human-friendly interface with an HTML form or by a RESTFull API, using GET or POST HTTP request methods, for single sequence or fasta file submission, respectively. The system is available at <http://portoreports.com/stm>.

## 6 Acknowledgments

Porto Reports is hosted at Free Web Hosting Area <http://freewebhostingarea.com>. This work was supported by Conselho Nacional de Desenvolvimento Científico e Tecnológico (CNPq), Coordenação de Aperfeiçoamento de Pessoal de Nível Superior (CAPES), Fundação de Apoio à Pesquisa do Distrito Federal (FAPDF) and Fundação de Apoio ao Desenvolvimento do Ensino, Ciência e Tecnologia do Estado de Mato Grosso do Sul (FUNDECT).

## References

1. Fjell, C. D., Hiss, J. A., Hancock, R. E. W. & Schneider, G. Designing antimicrobial peptides: form follows function. Nat. Rev. Drug Discov. 11, 37–51 (2011).

2. Porto, W. F., Silva, O. N. & Franco, O. L. Prediction and Rational Design of Antimicrobial Peptides. in Protein Structure (ed. Faraggi, E.) 377–396 (InTech, 2012). doi:10.5772/2335.

3. Franco, O. L. Elucidating Novel Bacterial Targets and Designing Unusual Antimicrobial Peptides: Two Faces of the Same Proteomic Coin. J. Proteomics Bioinform. s8, (2014).

4. Pinto, M. F. S. et al. Cyclotides: From Gene Structure to Promiscuous Multifunctionality. J. Evid. Based. Complementary Altern. Med. 17, 40–53 (2011).

5. Cândido, E. S. et al. Structural and functional insights into plant bactericidal peptides. in Science against Microbial Pathogens: Communicating Current Research and Technological Advances (ed. Méndez-Vilas, A.) 951–960 (Formatex, 2011).

6. Ribeiro, S. M. et al. Plant Antifungal Peptides. in Handbook of Biologically Active Peptides (ed. Kastin, A. J.) 169–179 (Elsevier Inc., 2013).

7. Silva, O. N. et al. Exploring the pharmacological potential of promiscuous host-defense peptides: from natural screenings to biotechnological applications. Front. Microbiol. 2, 232 (2011).

8. Porto, W. F., Pires, Á. S. & Franco, O. L. CS-AMPPred: an updated SVM model for antimicrobial activity prediction in cysteine-stabilized peptides. PLoS One 7, e51444 (2012).

9. Brand, G. D. et al. Probing protein sequences as sources for encrypted antimicrobial peptides. PLoS One 7, e45848 (2012).

10. Okubo, B. M. et al. Evaluation of an antimicrobial L-amino acid oxidase and peptide derivatives from Bothropoides mattogrosensis pitviper venom. PLoS One 7, e33639 (2012).

11. Mandal, S. M. et al. Identification of multifunctional peptides from human milk. Peptides 56, 84–93 (2014).

12. Loose, C., Jensen, K., Rigoutsos, I. & Stephanopoulos, G. A linguistic model for the rational design of antimicrobial peptides. Nature 443, 867–9 (2006).

13. Cherkasov, A. et al. Use of artificial intelligence in the design of small peptide antibiotics effective against a broad spectrum of highly antibiotic-resistant superbugs. ACS Chem. Biol. 4, 65–74 (2009).

14. Chou, K.-C. Some remarks on protein attribute prediction and pseudo amino acid composition. J. Theor. Biol. 273, 236–47 (2011).

15. Torrent, M., Andreu, D., Nogués, V. M. & Boix, E. Connecting peptide physicochemical and antimicrobial properties by a rational prediction model. PLoS One 6, e16968 (2011).

16. Waghu, F. H. et al. CAMP: Collection of sequences and structures of antimicrobial peptides. Nucleic Acids Res. 42, D1154–8 (2014).

17. Porto, W. F., Pires, Á. S. & Franco, O. L. Antimicrobial activity predictors benchmarking analysis using shuffled and designed synthetic peptides. J. Theor. Biol. 426, 96–103 (2017).

18. Eisenberg, D., Weiss, R. M., Terwilliger, T. C. & Wilcox, W. Hydrophobic moments and protein structure. Faraday Symp. Chem. Soc. 17, 109 (1982).

19. Porto, W. F., Fernandes, F. C. & Franco, O. L. An SVM Model Based on Physicochemical Properties to Predict Antimicrobial Activity from Protein Sequences with Cysteine Knot Motifs. in Advances in Bioinformatics and Computational Biology (eds. Ferreira, C. E., Miyano, S. & Stadler, P. F.) vol. 6268 59–62 (Springer Berlin Heidelberg, 2010).

20. Porto, W. F. et al. In silico optimization of a guava antimicrobial peptide enables combinatorial exploration for peptide design. Nat. Commun. 9, 1490 (2018).

21. Fjell, C. D. et al. Identification of novel antibacterial peptides by chemoinformatics and machine learning. J. Med. Chem. 52, 2006–15 (2009).

22. Lata, S., Mishra, N. K. & Raghava, G. P. S. AntiBP2: improved version of antibacterial peptide prediction. BMC Bioinformatics 11 Suppl 1, S19 (2010).

23. Fernandes, F. C., Rigden, D. J. & Franco, O. L. Prediction of antimicrobial peptides based on the adaptive neuro-fuzzy inference system application. Biopolymers 98, 280–7 (2012).

24. Porto, W. F., Pires, A. S. & Franco, O. L. Computational tools for exploring sequence databases as a resource for antimicrobial peptides. Biotechnol. Adv. 35, (2017).

25. Chou, K. C. Prediction of protein cellular attributes using pseudo-amino acid composition. Proteins 43, 246–55 (2001).

26. Xiao, X., Wang, P., Lin, W.-Z., Jia, J.-H. & Chou, K.-C. iAMP-2L: a two-level multi-label classifier for identifying antimicrobial peptides and their functional types. Anal. Biochem. 436, 168–77 (2013).

27. Veltri, D., Kamath, U. & Shehu, A. Deep learning improves antimicrobial peptide recognition. Bioinformatics 34, 2740–2747 (2018).

28. Yan, J. et al. Deep-AmPEP30: Improve Short Antimicrobial Peptides Prediction with Deep Learning. Mol. Ther. - Nucleic Acids 20, 882–894 (2020).

29. Porto, W. F., Fensterseifer, I. C. M., Ribeiro, S. M. & Franco, O. L. Joker: An algorithm to insert patterns into sequences for designing antimicrobial peptides. Biochim. Biophys. Acta - Gen. Subj. 1862, (2018).

30. Simm, S., Einloft, J., Mirus, O. & Schleiff, E. 50 years of amino acid hydrophobicity scales: revisiting the capacity for peptide classification. Biol. Res. 49, 31 (2016).

31. MacCallum, J. L. & Tieleman, D. P. Hydrophobicity scales: a thermodynamic looking glass into lipid–protein interactions. Trends Biochem. Sci. 36, 653–662 (2011).

32. Peters, C. & Elofsson, A. Why is the biological hydrophobicity scale more accurate than earlier experimental hydrophobicity scales? Proteins Struct. Funct. Bioinforma. 82, 2190–2198 (2014).

33. Zamora, W. J., Campanera, J. M. & Luque, F. J. Development of a Structure-Based, pH-Dependent Lipophilicity Scale of Amino Acids from Continuum Solvation Calculations. J. Phys. Chem. Lett. 10, 883–889 (2019).

34. Kyte, J. & Doolittle, R. F. A simple method for displaying the hydropathic character of a protein. J. Mol. Biol. 157, 105–32 (1982).

35. Radzicka, A. & Wolfenden, R. Comparing the Polarities of the Amino Acids: Side-Chain Distribution Coefficients between the Vapor Phase, Cyclohexane, 1 - 0ctano1, and Neutral Aqueous Solutiont. 1664–1670 (1988).

36. Boman, H. G. Antibacterial peptides: basic facts and emerging concepts. 197–215 (2003).

37. Nagarajan, D., Nagarajan, T., Nanajkar, N. & Chandra, N. A Uniform In Vitro Efficacy Dataset to Guide Antimicrobial Peptide Design. Data 4, 27 (2019).

38. Wimley, W. C. & White, S. H. Experimentally determined hydrophobicity scale for proteins at membrane interfaces. Nat. Struct. Mol. Biol. 3, 842–848 (1996).

39. Wang, G., Li, X. & Wang, Z. APD2: the updated antimicrobial peptide database and its application in peptide design. Nucleic Acids Res. 37, D933–7 (2009).

40. Jones, D. T. Protein secondary structure prediction based on position-specific scoring matrices 1 1Edited by G. Von Heijne. J. Mol. Biol. 292, 195–202 (1999).

41. Li, W. & Godzik, A. Cd-hit: a fast program for clustering and comparing large sets of protein or nucleotide sequences. Bioinformatics 22, 1658–9 (2006).

42. Glas, A. S., Lijmer, J. G., Prins, M. H., Bonsel, G. J. & Bossuyt, P. M. M. The diagnostic odds ratio: a single indicator of test performance. J. Clin. Epidemiol. 56, 1129–1135 (2003).

43. Vishnepolsky, B. & Pirtskhalava, M. Prediction of Linear Cationic Antimicrobial Peptides Based on Characteristics Responsible for Their Interaction with the Membranes. J. Chem. Inf. Model. 54, 1512–1523 (2014).

