## Supplementary Material for "Sense the Moment: a highly sensitive antimicrobial activity predictor based on hydrophobic moment"

Supplementary information

**Table S1** – The training data set.

| **APD ID** | **Sequence** | **GHM** | **Shuffled GHM** |
| --- | --- | --- | --- |
| AP00001 | GLWSKIKEVGKEAAKAAAKA AGKAALGAVSEAV | 0.540 | 0.291 |
| AP00011 | WNPFKELERAGQRVRDAVIS AAPAVATVGQAAAIARG | 0.650 | 0.347 |
| AP00013 | GLFDIIKKIAESF | 1.420 | 0.571 |
| AP00014 | GLLDIVKKVVGAFGSL | 1.099 | 0.452 |
| AP00017 | GLFDIVKKVVGTLAGL | 1.045 | 0.452 |
| AP00019 | GLFDIAKKVIGVIGSL | 1.098 | 0.458 |
| AP00020 | GLFDIVKKIAGHIAGSI | 1.033 | 0.447 |
| AP00035 | KSSAYSLQMGATAIKQVKKL FKKWGW | 0.796 | 0.336 |
| AP00050 | GIGASILSAGKSALKGLAKG LAEHFAN | 0.679 | 0.308 |
| AP00055 | IIGPVLGMVGSALGGLLKKI | 0.897 | 0.344 |
| AP00058 | GIGTKILGGVKTALKGALKE LASTYAN | 0.702 | 0.319 |
| AP00064 | ILGPVLGLVGNALGGLIKNE | 0.686 | 0.376 |
| AP00066 | IKITTMLAKLGKVLAHV | 0.915 | 0.445 |
| AP00069 | INIKDILAKLVKVLGHV | 1.092 | 0.515 |
| AP00070 | INVLGILGLLGKALSHL | 0.707 | 0.407 |
| AP00071 | FLPAIFRMAAKVVPTIICSI TKKC | 0.908 | 0.39 |
| AP00073 | FLPLLAGLAANFLPKIFCKI TRKC | 0.767 | 0.397 |
| AP00074 | FLPVLAGIAAKVVPALFCKI TKKC | 0.549 | 0.338 |
| AP00076 | GILDTLKNLAISAAKGAAQG LVNKASCKLSGQC | 0.365 | 0.297 |
| AP00078 | GILLDKLKNFAKTAGKGVLQ SLLNTASCKLSGQC | 0.451 | 0.303 |
| AP00083 | GILSLVKGVAKLAGKGLAKE GGKFGLELIACKIAKQC | 0.276 | 0.293 |
| AP00085 | SLFSLIKAGAKFLGKNLLKQ GACYAACKASKQC | 0.198 | 0.291 |
| AP00086 | GIMSIVKDVAKNAAKEAAKG ALSTLSCKLAKTC | 0.333 | 0.3 |
| AP00088 | GILDTLKQFAKGVGKDLVKG AAQGVLSTVSCKLAKTC | 0.365 | 0.29 |
| AP00089 | FLGALFKVASKVLPSVFCAI TKKC | 0.780 | 0.344 |
| AP00095 | LLPIVGNLLKSLL | 1.158 | 0.488 |
| AP00097 | VLPIIGNLLNSLL | 1.156 | 0.476 |
| AP00101 | FVQWFSKFLGRIL | 1.366 | 0.612 |
| AP00105 | FLPLFASLIGKLL | 0.939 | 0.425 |
| AP00106 | FLPFLASLLTKVL | 0.997 | 0.428 |
| AP00109 | VLPLISMALGKLL | 0.796 | 0.424 |
| AP00110 | NFLGTLINLAKKIM | 1.250 | 0.52 |
| AP00111 | FLPILINLIHKGLL | 0.838 | 0.488 |
| AP00117 | FLPFIARLAAKVFPSIICSV TKKC | 0.821 | 0.389 |
| AP00119 | GILSSIKGVAKGVAKNVAAQ LLDTLKCKITGC | 0.479 | 0.314 |
| AP00123 | GLFLDTLKGLAGKLLQGLKC IKAGCKP | 0.770 | 0.345 |
| AP00126 | GGLKKLGKKLEGVGKRVFKA SEKALPVAVGIKALGK | 0.767 | 0.333 |
| AP00127 | RWKVFKKIEKVGRNIRDGVI KAAPAIEVLGQAKAL | 0.708 | 0.385 |
| AP00129 | GWLKKIGKKIERVGQNTRDA TVKGLEVAQQAANVAATVR | 0.874 | 0.337 |
| AP00134 | SWLSKTAKKLENSAKKRISE GIAIAIQGGPR | 0.750 | 0.359 |
| AP00136 | FLPLILRKIVTAL | 1.113 | 0.618 |
| AP00140 | SQLGDLGSGAGQGGGGGGSI RAAGGAFGKLEAAREEEFFY KKQKEQLERLKNDQIHQAEF HHQQIKEHEEAIQRHKDFLN NLHK | 0.409 | 0.205 |
| AP00142 | GLKKLLGKLLKKLGKLLLK | 1.191 | 0.485 |
| AP00144 | GIGKFLHSAKKFGKAFVGEI MNS | 0.922 | 0.368 |
| AP00146 | GIGAVLKVLTTGLPALISWI KRKRQQ | 0.793 | 0.42 |
| AP00155 | RGLRRLGRKIAHGVKKYGPT VLRIIRIAG | 1.529 | 0.473 |
| AP00157 | ALWKTMLKKLGTMALHAGKA ALGAAADTISQGTQ | 0.598 | 0.265 |
| AP00159 | ALWKNMLKGIGKLAGKAALG AVKKLVGAES | 0.584 | 0.319 |
| AP00164 | ALWKTIIKGAGKMIGSLAKN LLGSQAQPES | 0.618 | 0.311 |
| AP00166 | GWGSFFKKAAHVGKHVGKAA LTHYL | 0.658 | 0.314 |
| AP00200 | LKLKSIVSWAKKVL | 0.890 | 0.542 |
| AP00201 | INLKALAALAKKIL | 0.662 | 0.534 |
| AP00209 | GVLSNVIGYLKKLGTGALNA VLKQ | 0.831 | 0.362 |
| AP00233 | GWIRDFGKRIERVGQHTRDA TIQTIAVAQQAANVAATLKG | 0.786 | 0.34 |
| AP00240 | GLLSVLGSVAKHVLPHVVPV IAEHL | 0.695 | 0.321 |
| AP00249 | GLVSSIGRALGGLLADVVKS KGQPA | 0.793 | 0.375 |
| AP00254 | GLWEKIKEKASELVSGIVEG VK | 1.091 | 0.409 |
| AP00257 | GLWQKIKSAAGDLASGIVEG IKS | 0.997 | 0.367 |
| AP00260 | GLFVGVLAKVAAHVVPAIAE HF | 0.668 | 0.321 |
| AP00262 | GFVDFLKKVAGTIANVVT | 1.159 | 0.423 |
| AP00263 | GLLQTIKEKLESLESLAKGI VSGIQA | 0.991 | 0.364 |
| AP00278 | VFHLLGKIIHHVGNFVYGFS HVF | 1.140 | 0.348 |
| AP00315 | SLGSFLKGVGTTLASVGKVV SDQFGKLLQAGQG | 0.902 | 0.283 |
| AP00316 | GIVDFAKKVVGGIRNALGI | 1.082 | 0.492 |
| AP00322 | GILDVAKTLVGKLRNVLGI | 1.029 | 0.5 |
| AP00323 | GVLDAFRKIATVVKNVV | 1.384 | 0.543 |
| AP00324 | GVGDLIRKAVSVIKNIV | 1.452 | 0.556 |
| AP00325 | GVIDAAKKVVNVLKNLF | 1.330 | 0.505 |
| AP00326 | GVGSFIHKVVSAIKNVA | 1.182 | 0.432 |
| AP00330 | GWLRKAAKSVGKFYYKHKYY IKAAWQIGKHAL | 0.884 | 0.327 |
| AP00339 | FFGWLIKGAIHAGKAIHGLI HRRRH | 0.687 | 0.458 |
| AP00347 | RWKFFKKIERVGQNVRDGLI KAGPAIQVLGAAKAL | 0.565 | 0.375 |
| AP00350 | PWNIFKEIERAVARTRDAVI SAGPAVRTVAAATSVAS | 0.648 | 0.343 |
| AP00358 | FGLPMLSILPKALCILLKRK C | 0.470 | 0.438 |
| AP00366 | GRFKRFRKKFKKLFKKLSPV IPLLHLG | 0.819 | 0.46 |
| AP00368 | GLFRRLRDSIRRGQQKILEK ARRIGERIKDIFRG | 1.514 | 0.457 |
| AP00371 | GLLSRLRDFLSDRGRRLGEK IERIGQKIKDLSEFFQS | 1.183 | 0.397 |
| AP00373 | AKIPIKAIKTVGKAVGKGLR AINIASTANDVFNFLKPKKR KH | 0.386 | 0.322 |
| AP00374 | GKVWDWIKSTAKKLWNSEPV KELKNTALNAAKNLVAEKIG ATPS | 0.249 | 0.268 |
| AP00376 | GWKDWAKKAGGWLKKKGPGM AKAALKAAMQ | 0.648 | 0.296 |
| AP00377 | GWKDWLKKGKEWLKAKGPGI VKAALQAATQ | 0.686 | 0.32 |
| AP00379 | DFKDWMKTAGEWLKKKGPGI LKAAMAAAT | 0.823 | 0.315 |
| AP00382 | GLVDVLGKVGGLIKKLLPG | 1.053 | 0.417 |
| AP00383 | LLKELWTKMKGAGKAVLGKI KGLL | 0.882 | 0.382 |
| AP00385 | FKLGSFLKKAWKSKLAKKLR AKGKEMLKDYAKGLLEGGSE EVPGQ | 0.153 | 0.28 |
| AP00386 | WLGSALKIGAKLLPSVVGLF KKKKQ | 0.788 | 0.367 |
| AP00388 | GIWGTLAKIGIKAVPRVISM LKKKKQ | 0.821 | 0.409 |
| AP00389 | GIWGTALKWGVKLLPKLVGM AQTKKQ | 0.630 | 0.338 |
| AP00390 | FWGALIKGAAKLIPSVVGLF KKKQ | 0.727 | 0.371 |
| AP00391 | FIGTALGIASAIPAIVKLFK | 0.646 | 0.345 |
| AP00399 | HVDKKVADKVLLLKQLRIMR LLTRL | 0.412 | 0.508 |
| AP00405 | FISAIASMLGKFL | 1.035 | 0.416 |
| AP00408 | FLFPLITSFLSKVL | 0.854 | 0.423 |
| AP00414 | SIGSALKKALPVAKKIGKIA LPIAKAALP | 0.931 | 0.322 |
| AP00417 | SIGTAVKKAVPIAKKVGKVA IPIAKAVLSVVGQLVG | 0.962 | 0.285 |
| AP00418 | GLRKRLRKFRNKIKEKLKKI GQKIQGFVPKLAPRTDY | 0.904 | 0.383 |
| AP00424 | GFLGPLLKLAAKGVAKVIPH LIPSRQQ | 0.652 | 0.369 |
| AP00427 | GLLGPLLKIAAKVGSNLL | 0.924 | 0.401 |
| AP00430 | ILGKIWEGIKSLF | 1.262 | 0.529 |
| AP00433 | SSLLEKGLDGAKKAVGGLGK LGKDAVEDLESVGKGAVHDV KDVLDSV | 1.142 | 0.269 |
| AP00434 | GLMSVLGHAVGNVLGGLFKS | 0.790 | 0.334 |
| AP00435 | GWFGKAFRSVSNFYKKHKTY IHAGLSAATLL | 0.743 | 0.317 |
| AP00446 | GADFQECMKEHSQKQHQHQG | 0.364 | 0.323 |
| AP00453 | FLPAIVGAAGQFLPKIFCAI SKKC | 0.749 | 0.326 |
| AP00455 | FFPIVAGVAGQVLKKIYCTI SKKC | 0.669 | 0.356 |
| AP00459 | FITLLLRKFICSITKKC | 0.783 | 0.531 |
| AP00461 | FLPMLAGLAASMVPKLVCLI TKKC | 0.604 | 0.313 |
| AP00470 | FLPIIASVAAKVFSKIFCAI SKKC | 0.716 | 0.347 |
| AP00474 | FIHHIFRGIVHAGRSIGRFL TG | 1.479 | 0.491 |
| AP00475 | GLNTLKKVFQGLHEAIKLIN NHVQ | 1.229 | 0.405 |
| AP00485 | GFGALFKFLAKKVAKTVAKQ AAKQGAKYVVNKQME | 0.078 | 0.305 |
| AP00492 | RQRVEELSKFSKKGAAARRR K | 0.593 | 0.511 |
| AP00493 | NLVSGLIEARKYLEQLHRKL KNCKV | 0.800 | 0.453 |
| AP00494 | KWKLFKKIPKFLHLAKKF | 0.963 | 0.491 |
| AP00496 | AKKVFKRLEKLFSKIQNDK | 1.166 | 0.515 |
| AP00497 | ILGPVLGLVSDTLDDVLGIL | 0.959 | 0.388 |
| AP00498 | GLVRKGGEKFGEKLRKIGQK IKEFFQKLALEIEQ | 1.187 | 0.374 |
| AP00499 | VGALAVVVWLWLWLW | 0.209 | 0.178 |
| AP00501 | GIGKHVGKALKGLKGLLKGL GES | 1.100 | 0.375 |
| AP00502 | FLRFIGSVIHGIGHLVHHIG VAL | 1.176 | 0.392 |
| AP00503 | FLGVVFKLASKVFPAVFGKV | 0.766 | 0.367 |
| AP00506 | KLAKLAKKLAKLAK | 1.328 | 0.56 |
| AP00510 | ILQKAVLDCLKAAGSSLSKA AITAIYNKIT | 0.374 | 0.323 |
| AP00514 | FLGGLMKAFPALICAVTKKC | 0.748 | 0.35 |
| AP00516 | IWLTALKFLGKHAAKHLAKQ QLSKL | 0.231 | 0.374 |
| AP00517 | KIKWFKTMKSIAKFIAKEQM KKHLGGE | 0.295 | 0.366 |
| AP00522 | INWLKLGKAIIDAL | 0.880 | 0.542 |
| AP00527 | ILGPVISKIGGVLGGLLKNL | 0.983 | 0.381 |
| AP00533 | GVVDILKGAAKDIAGHLASK VMNKL | 0.520 | 0.378 |
| AP00535 | GLGSVFGRLARILGRVIPKV AKKLGPKVAKVLPKVMKEAI PMAVEMAKSQEEQQPQ | 0.684 | 0.271 |
| AP00540 | GLLDTLKGAAKNVVGSLASK VMEKL | 0.612 | 0.363 |
| AP00541 | IDWKKLLDAAKQIL | 0.949 | 0.583 |
| AP00544 | GVLDIFKDAAKQILAHAAEK QI | 0.569 | 0.423 |
| AP00546 | FLSLIPHAINAVSAIAKHN | 0.835 | 0.399 |
| AP00556 | GFMKYIGPLIPHAVKAISDL I | 1.158 | 0.372 |
| AP00557 | RVKRVWPLVIRTVIAGYNLY RAIKKK | 0.863 | 0.486 |
| AP00569 | FLPLLLAGLPLKLCFLFKKC | 0.403 | 0.366 |
| AP00572 | GIMDTIKDTAKTVAVGLLNK LKCKITGC | 0.623 | 0.352 |
| AP00575 | GLLDTFKNLALNAAKSAGVS VLNSLSCKLSKTC | 0.324 | 0.296 |
| AP00577 | GLFTLIKGAAKLIGKTVAKE AGKTGLELMACKITNQC | 0.212 | 0.284 |
| AP00582 | GFSSLFKAGAKYLLKSVGKA GAQQLACKAANNCA | 0.213 | 0.275 |
| AP00583 | GVITDALKGAAKTVAAELLR KAHCKLTNSC | 0.335 | 0.344 |
| AP00584 | VIDDLKKVAKKVRRELLCKK HHKKLN | 0.623 | 0.458 |
| AP00586 | FLPLLFGAISHLL | 0.845 | 0.37 |
| AP00588 | FLGSIVGALASALPSLISKI RN | 0.986 | 0.401 |
| AP00591 | FLPLLGNLLRGLL | 1.253 | 0.595 |
| AP00598 | FLSAITSLLGKLL | 1.037 | 0.441 |
| AP00599 | GIWDTIKSMGKVFAGKILQN L | 0.685 | 0.406 |
| AP00601 | FLSLALAALPKFLCLVFKKC | 0.661 | 0.363 |
| AP00605 | ILPILSLIGGLLGK | 0.849 | 0.411 |
| AP00610 | GLFSVVTGVLKAVGKNVAKN VGGSLLEQLKCKISGGC | 0.254 | 0.282 |
| AP00611 | FIGPIISALASLFG | 0.858 | 0.317 |
| AP00614 | ALFSILRGLKKLGKMGQAFV NCEIYKKC | 0.640 | 0.379 |
| AP00624 | ALLGDFFRKSKEKIGKEFKR IVQRIKDFLRNLVPRTES | 1.134 | 0.387 |
| AP00639 | GLIGSIGKALGGLLVDVLKP KLQAAS | 0.694 | 0.337 |
| AP00640 | GLLGLLGSVVSHVVPAIVGH F | 0.652 | 0.286 |
| AP00651 | GLFSILKGVGKIALKGLAKN MGKMGLDLVSCKISKEC | 0.222 | 0.302 |
| AP00654 | GLLDTIKNTAKNLAVGLLDK IKCKMTGC | 0.694 | 0.356 |
| AP00656 | GIMDSVKNAAKNLAGQLLDT IKCKITAC | 0.804 | 0.345 |
| AP00660 | FWGALAKGALKLIPSLFSSF SKKD | 0.670 | 0.351 |
| AP00661 | GILSLFTGGIKALGKTLFKM AGKAGAEHLACKATNQC | 0.307 | 0.266 |
| AP00664 | FLPAIAGILSQLF | 0.934 | 0.385 |
| AP00677 | GLRKKFRKTRKRIQKLGRKI GKTGRKVWKAWREYGQIPYP CRI | 0.984 | 0.353 |
| AP00678 | RLKELITTGGQKIGEKIRRI GQRIKDFFKNLQPREEKS | 1.186 | 0.375 |
| AP00686 | KRFGRLAKSFLRMRILLPRR KILLAS | 0.730 | 0.524 |
| AP00691 | GFFKKAWRKVKHAGRRVLDT AKGVGRHYVNNWLNRYR | 1.045 | 0.369 |
| AP00694 | AIGSILGALAKGLPTLISWI KNR | 0.874 | 0.401 |
| AP00696 | GLFDIIKNIVSTL | 1.352 | 0.553 |
| AP00699 | GLWQFIKDKLKDAATGLVTG IQS | 0.918 | 0.385 |
| AP00701 | GLLGSIGNAIGAFIANKLKP | 0.638 | 0.381 |
| AP00704 | GLLGSIGKVLGGYLAEKLKP K | 0.433 | 0.385 |
| AP00722 | GLLNGLALRLGKRALKKIIK RLCR | 0.943 | 0.522 |
| AP00723 | SLLSLIRKLIT | 1.385 | 0.723 |
| AP00736 | RLGDILQKAREKIEGGLKKL VQKIKDFFGKFAPRTES | 1.033 | 0.364 |
| AP00738 | GLVTGLLKTAGKLLGDLFGS LTG | 0.870 | 0.326 |
| AP00753 | VQETQKLAKTVGANLEETNK KLAPQIKSAYDDFVKQAQEV QKKLHEAASKQ | 0.876 | 0.249 |
| AP00754 | ETESTPDYLKNIQQQLEEYT KNFNTQVQNAFDSDKIKSEV NNFIESLGKILNTEKKEAPK | 0.419 | 0.227 |
| AP00755 | ENFFKEIERAGQRIRDAIIS AAPAVETLAQAQKIIKGGD | 0.715 | 0.344 |
| AP00756 | ALWKDILKNAGKAALNEINQ LVNQ | 0.901 | 0.407 |
| AP00764 | GLRSKIWLWVLLMIWQESNK FKKM | 0.267 | 0.425 |
| AP00769 | GLLGAMFKVASKVLPHVVPA ITEHF | 0.814 | 0.317 |
| AP00772 | FRGLAKLLKIGLKSFARVLK KVLPKAAKAGKALAKSMADE NAIRQQNQ | 0.734 | 0.307 |
| AP00773 | GKFSVFGKILRSIAKVFKGV GKVRKQFKTASDLDKNQ | 0.650 | 0.346 |
| AP00782 | GWGSIFKHGRHAAKHIGHAA VNHYL | 0.678 | 0.361 |
| AP00786 | GWKKWFNRAKKVGKTVGGLA VDHYL | 0.705 | 0.393 |
| AP00788 | AGWGSIFKHIFKAGKFIHGA IQAHND | 0.748 | 0.346 |
| AP00791 | GWKKWLRKGAKHLGQAAIKG LAS | 0.818 | 0.402 |
| AP00792 | FLGLLFHGVHHVGKWIHGLI HGHH | 0.867 | 0.335 |
| AP00817 | FLPLLASLFSRLL | 1.099 | 0.563 |
| AP00818 | FLPLIGKILGTIL | 1.049 | 0.439 |
| AP00822 | GIFNVFKGALKTAGKHVAGS LLNQLKCKVSGEC | 0.346 | 0.308 |
| AP00823 | SILPTIVSFLSKVF | 1.139 | 0.438 |
| AP00867 | FLPVIAGLLSKLF | 0.924 | 0.426 |
| AP00869 | ILPLVGNLLNDLL | 1.227 | 0.523 |
| AP00871 | FLPFLKSILGKIL | 1.010 | 0.505 |
| AP00872 | FLPFFASLLGKLL | 0.927 | 0.413 |
| AP00875 | FLSSIGKILGNLL | 1.245 | 0.489 |
| AP00876 | FLSIIAKVLGSLF | 1.038 | 0.444 |
| AP00877 | FLGSLIGAAIPAIKQLLGLK K | 0.641 | 0.38 |
| AP00879 | GRLRNLIEKAGQNIRGKIQG IGRRIKDILKNLQPRPQV | 1.084 | 0.397 |
| AP00883 | FLPLIASVAANLAPKIICKI TKTC | 0.732 | 0.343 |
| AP00893 | DVKGMKKAIKGILDCVIEKG YDKLAAKLKKVIQQLWE | 0.537 | 0.334 |
| AP00894 | GLLDFVTGVGKDIFAQLIKQ I | 0.829 | 0.421 |
| AP00900 | FLSHIAGFLSNLF | 1.061 | 0.432 |
| AP00901 | GWMSKIASGIGTFLSGVQQG | 0.896 | 0.331 |
| AP00911 | FLSLIPHIVSGVAALAKHL | 1.021 | 0.359 |
| AP00939 | ALWKTLLKGAGKVFGHVAKQ FLGSQGQPES | 0.618 | 0.3 |
| AP00940 | GLWSKIKEAAKTAGKMAMGF VNDMV | 0.911 | 0.344 |
| AP00942 | GLWKSLLKNVGVAAGKAALN AVTDMVNQ | 0.448 | 0.329 |
| AP00949 | GLWSTIKQKGKEAAIAAAKA AGQAVLNSASEAL | 0.387 | 0.287 |
| AP00959 | ALWKTLLKKVGKVAGKAVLN AVTNMANQNEQ | 0.595 | 0.332 |
| AP00960 | AVWKDFLKNIGKAAGKAVLN SVTDMVNE | 0.637 | 0.356 |
| AP00964 | GLWSKIKEAAKAAGKAALNA VTGLVNQGDQPS | 0.815 | 0.299 |
| AP00965 | SVLSTITDMAKAAGRAALNA ITGLVNQ | 1.022 | 0.354 |
| AP00973 | LLGMIPLAISAISALSKL | 0.677 | 0.343 |
| AP00974 | FLSLLPSLVSGAVSLVKIL | 0.916 | 0.344 |
| AP01011 | GLFGKLIKKFGRKAISYAVK KARGKH | 0.744 | 0.424 |
| AP01012 | SWKSMAKKLKEYMEKLKQRA | 1.017 | 0.432 |
| AP01014 | GLKDKFKSMGEKLKQYIQTW KAKF | 0.856 | 0.375 |
| AP01016 | GFFGKMKEYFKKFGASFKRR FANLKKRL | 0.936 | 0.414 |
| AP01019 | GETFDKLKEKLKTFYQKLVE KAEDLKGDLKAKLS | 0.880 | 0.328 |
| AP01129 | GLGSLVGNALRIGAKLL | 1.119 | 0.484 |
| AP01130 | GMASKAGSVLGKVAKVALKA AL | 0.786 | 0.347 |
| AP01158 | ALYKKFKKKLLKSLKRL | 1.105 | 0.556 |
| AP01217 | GFRDVLKGAAKQFVKTVAGH IANI | 0.682 | 0.415 |
| AP01219 | GFKDWIKGAAKKLIKTVAAN IANQ | 0.540 | 0.391 |
| AP01223 | GFKDLLKGAAKALVKTVLF | 0.811 | 0.431 |
| AP01236 | GLLDFAKHVIGIASKL | 1.075 | 0.464 |
| AP01240 | ALKAALLAILKIVRVIKK | 1.119 | 0.539 |
| AP01241 | FASLLGKALKALAKQ | 1.058 | 0.466 |
| AP01242 | GLLSFLPKVIGVIGHLIHP PS | 0.888 | 0.34 |
| AP01248 | INMKASAAVAKKLL | 0.361 | 0.509 |
| AP01249 | GILDAIKAIAKAAG | 1.105 | 0.478 |
| AP01258 | GLMDVFKGAAKNLLASALD KIRCKVTKC | 0.573 | 0.383 |
| AP01260 | IIGHLIKTALGMLGL | 0.749 | 0.402 |
| AP01261 | IIEKLVNTALGLLSGL | 1.082 | 0.445 |
| AP01262 | GLADFLNKAVGKVVDFVKS | 1.215 | 0.448 |
| AP01263 | FLPLVTMLLGKLF | 0.774 | 0.41 |
| AP01264 | RIGVLLARLPKLFSLFKLMG KKV | 0.654 | 0.476 |
| AP01266 | AVDLAKIANKVLSSLF | 1.058 | 0.476 |
| AP01269 | GFLSILKKVLPKVMAHMK | 0.901 | 0.439 |
| AP01292 | GVLGAVKDLLIGAGKSAAQS VLKTLSCKLSNDC | 0.313 | 0.302 |
| AP01303 | VIPFVASVAAEMMQHVYCAA SKKC | 0.614 | 0.314 |
| AP01328 | GFIFHIIKGLFHAGKMIHGL V | 0.927 | 0.366 |
| AP01331 | IFGAILPLALGALKNLIK | 0.869 | 0.398 |
| AP01332 | FIGAILPAIAGLVHGLINR | 0.830 | 0.44 |
| AP01345 | FFGTALKIAANVLPTAICKI LKKC | 0.778 | 0.356 |
| AP01346 | FFPLVLGALGSILPKIF | 0.784 | 0.341 |
| AP01347 | FIITGLVRGLTKLF | 1.217 | 0.572 |
| AP01353 | FWGHIWNAVKRVGANALHGA VTGALS | 0.708 | 0.349 |
| AP01354 | GFWKKVGSAAWGGVKAAAKG AAVGGLNALAKHIQ | 0.278 | 0.268 |
| AP01383 | GLVSGLLNTAGGLLGDLLGS LGSLSGGES | 0.524 | 0.26 |
| AP01384 | GMWGSLLKGVATVVKHVLPH ALSSQQS | 0.714 | 0.303 |
| AP01386 | LLGDLLGQTSKLVNDLTDTV GSIV | 1.100 | 0.374 |
| AP01388 | GLLSGILNSAGGLLGNLIGS LSN | 0.836 | 0.318 |
| AP01395 | GILLNTLKGAAKNVAGVLLD KLKCKITGGC | 0.489 | 0.331 |
| AP01396 | GLMDSLKGLAATAGKTVLQG LLKTASCKLEKTC | 0.385 | 0.288 |
| AP01397 | ILPFLAGLFSKIL | 0.951 | 0.431 |
| AP01402 | GVVDILKGAGKDLLAHALSK LSEKV | 0.498 | 0.382 |
| AP01423 | FLPAVLRVAAKIVPTVFCAI SKKC | 0.885 | 0.387 |
| AP01428 | GFKGAFKNVMFGIAKSAGKS ALNALACKIDKSC | 0.191 | 0.3 |
| AP01429 | GLLDSFKNAMIGIAKSAGKT ALNKIACKIDKTC | 0.385 | 0.318 |
| AP01432 | FMGGLIKAATKIVPAAYCAI TKKC | 0.687 | 0.324 |
| AP01434 | FFGSVLKLIPKIL | 1.252 | 0.506 |
| AP01443 | GLFLNTVKDVAKDVAKDVAG KLLESLKCKITGCKP | 0.635 | 0.321 |
| AP01445 | FMGSALRIAAKVLPAALCQI FKKC | 0.754 | 0.389 |
| AP01449 | FLGAIAAALPHVINAVTNAL | 0.951 | 0.323 |
| AP01452 | IIGAIAAALPHVINAIKNTF | 0.972 | 0.377 |
| AP01455 | FFPLALLCKVFKKC | 0.654 | 0.48 |
| AP01457 | GLKDIFKAGLGSLVKGIAAH VAN | 0.510 | 0.371 |
| AP01461 | ILGKLLSTAAGLLSNL | 0.992 | 0.403 |
| AP01462 | ILGAILPLVSGLLSNKL | 0.523 | 0.392 |
| AP01465 | VNWKKVLGKIIKVAK | 0.980 | 0.555 |
| AP01499 | FLPVLARLAVKFLPSIVCAA TKKC | 0.754 | 0.386 |
| AP01512 | FFSLLPSLIGGLVSAIK | 0.969 | 0.36 |
| AP01516 | LNLKGIFKKVASLLT | 0.885 | 0.502 |
| AP01517 | INLLKIAKGIIKSL | 1.116 | 0.557 |
| AP01541 | AVLDILKDVGKGLLSHFMEK V | 0.853 | 0.427 |
| AP01542 | AVLDFIKAAGKGLVTNIMEK VG | 0.664 | 0.394 |
| AP01544 | IFGAIAGLLKNIF | 1.035 | 0.463 |
| AP01545 | FFGHLFKLATKIIPSLFQ | 1.154 | 0.416 |
| AP01547 | GVIKSVLKGVAKTVALGML | 0.659 | 0.392 |
| AP01558 | LGAWLAGKVAGTVATYAWNR YV | 0.307 | 0.36 |
| AP01568 | DSHEKRHHEHRRKFHEKHHS HRGY | 0.296 | 0.322 |
| AP01579 | FVLPLVMCKILRKC | 0.701 | 0.593 |
| AP01583 | GWANTLKNVAGGLCKITGAA | 0.800 | 0.344 |
| AP01614 | WRSLGRTLLRLSHALKPLAR RSGW | 1.241 | 0.488 |
| AP01633 | RRWVRRVRRWVRRVVRVVRR WVRR | 2.228 | 0.64 |
| AP01634 | INWKKIFEKVKNLV | 1.049 | 0.602 |
| AP01637 | INWKKIASIGKEVLKAL | 0.928 | 0.497 |
| AP01638 | INWKKIAEVGGKILSSL | 0.880 | 0.473 |
| AP01641 | IDWLKLGKMVMDVL | 0.822 | 0.537 |
| AP01645 | GAFGNFLKGVAKKAGLKILS IAQCKLFGTC | 0.325 | 0.308 |
| AP01648 | GKLNLFLSRLEILKLFVGAL | 0.242 | 0.475 |
| AP01679 | GRILSFIKGLAEHL | 1.242 | 0.586 |
| AP01680 | ILGIITSLLKSLGKK | 1.071 | 0.496 |
| AP01681 | KDLHTVVSAILQAL | 1.200 | 0.504 |
| AP01699 | GRGREFMSNLKEKLSGVKEK MKNS | 0.841 | 0.412 |
| AP01700 | VKLIQIRIWIQYVTVLQMFS MKTKQ | 0.282 | 0.42 |
| AP01708 | SFLTTFKDLAIKAAKSAGQS VLSTLSCKLSNTC | 0.471 | 0.286 |
| AP01719 | GILDTFKGVAKGVAKDLAVH MLENLKCKMTGC | 0.407 | 0.32 |
| AP01739 | GIGGVLLGAGKATLKGLAKV LAEKYAN | 0.397 | 0.32 |
| AP01743 | GIGGALLSVGKLALKGLANV LADKFAN | 0.484 | 0.332 |
| AP01749 | LVQRGRFGRFLKKVRRFIPK VIIAAQIGSRFG | 0.665 | 0.451 |
| AP01753 | GIWSSIKNLASKAWNSDIGQ SLRNKAAGAINKFVADKIGV TPSQAASMTLDEIVDAMYYD | 0.222 | 0.229 |
| AP01768 | QLGELIQQGGQKIVEKIQKI GQRIRDFFSNLRPRQEA | 1.069 | 0.373 |
| AP01771 | ILPLLLGKVVCAITKKC | 0.559 | 0.432 |
| AP01778 | DSMGAVKLAKLLIDKMKCEV TKAC | 0.549 | 0.383 |
| AP01786 | GKLQAFLAKMKEIAAQTL | 0.649 | 0.44 |
| AP01794 | FVDLKKIANIINSIFGK | 0.915 | 0.512 |
| AP01824 | FLPKLFAKITKKNMAHIR | 0.779 | 0.518 |
| AP01851 | SFLSTFKELAINAAKNAGQS LLHTLSCKLDKTC | 0.559 | 0.3 |
| AP01887 | GFMDTAKNVAKNMAGNLLDN LKCKITKAC | 0.661 | 0.339 |
| AP01895 | FLGSLLGLVGKVVPTLFCKI SKKC | 0.810 | 0.347 |
| AP01896 | GLMSTLKDFGKTAAKEIAQS LLSTASCKLAKTC | 0.351 | 0.284 |
| AP01899 | FLKPLFNAALKLLP | 1.101 | 0.479 |
| AP01900 | FLPVLAGVLSRA | 0.968 | 0.588 |
| AP01901 | GLASFLGKALKAGLKIGSHL LGGAPQQ | 0.816 | 0.309 |
| AP01907 | GIGSALAKAAKLVAGIV | 0.912 | 0.367 |
| AP01920 | FTSKKSMLLFFFLGTISLSL CQ | 0.227 | 0.345 |
| AP01922 | GMWSKILGHLIR | 1.211 | 0.638 |
| AP01923 | GKWMSLLKHILK | 1.172 | 0.563 |
| AP01925 | GLLDAIKDTAQNLFANVLDK IKCKFTKC | 0.765 | 0.373 |
| AP01929 | GIFALIKTAAKFVGKNLLKQ AGKAGLEHLACKANNQC | 0.158 | 0.295 |
| AP01932 | FFPIVGKLLFGLSGLL | 0.798 | 0.357 |
| AP01943 | SLWETIKNAGKGFIQNLDKI R | 1.116 | 0.463 |
| AP01947 | FLGPIIKIATGILPTAICKF LKKC | 0.639 | 0.352 |
| AP01948 | SIRDKIKTIAIDLAKSAGTG VLKTLICKLDKSC | 0.248 | 0.354 |
| AP01953 | FALGAVTKLLPSLLCMITRK C | 0.792 | 0.413 |
| AP01956 | GFGMALKLLKKVL | 1.056 | 0.517 |
| AP01959 | AILTTLANWARKFL | 1.166 | 0.562 |
| AP01964 | IKLSPETKDNLKKVLKGAIK GAIAVAKMV | 0.911 | 0.363 |
| AP01966 | IKIPAVVKDTLKKVAKGVLS AVAGALTQ | 0.789 | 0.349 |
| AP01977 | FLFSLIPSAISGLISAFKGR R | 0.487 | 0.46 |
| AP01978 | FIGAIARLLSKIFGKR | 0.821 | 0.607 |
| AP02001 | GMATKAGTALGKVAKAVIGA AL | 0.705 | 0.312 |
| AP02002 | GLFLDTLKKFAKAGMEAVIN PK | 0.759 | 0.397 |
| AP02003 | GFWTTAAEGLKKFAKAGLAS ILNPK | 0.605 | 0.33 |
| AP02004 | GVWTTILGGLKKFAKGGLEA LTNPK | 0.818 | 0.335 |
| AP02006 | GLLDALSGILGL | 0.851 | 0.439 |
| AP02011 | GLFDVIKKVASVIGLASP | 0.891 | 0.41 |
| AP02013 | FIGKLISAASGLLSHL | 0.990 | 0.391 |
| AP02026 | GFWGKLWEGVKSAI | 0.956 | 0.449 |
| AP02038 | FLGLIFHGLVHAGKLIHGLI HRNRG | 0.725 | 0.431 |
| AP02059 | FLSTALKVAANVVPTLFCKI TKKC | 0.820 | 0.352 |
| AP02064 | GILDKLKEFGISAARGVAQS LLNTTASCKLAKTC | 0.599 | 0.315 |
| AP02103 | GLWDTIKQAGKKFFLNVLDK IRCKVAGGCRT | 0.709 | 0.381 |
| AP02104 | MQFITDLIKKAVDFFKGLFG NK | 0.990 | 0.425 |
| AP02105 | MAADIISTIGDLVKLIINTV KKFQK | 1.202 | 0.407 |
| AP02118 | FLPAALAGIGGILGKLF | 0.691 | 0.314 |
| AP02135 | FIHHIIGGLFSAGKAIHRLI RRRRR | 1.199 | 0.537 |
| AP02136 | FIHHIIGWISHGVRAIHRAI HG | 1.355 | 0.461 |
| AP02137 | FLHHIVGLIHHGLSLFGDRA D | 0.920 | 0.443 |
| AP02140 | IWDAIFHGAKHFLHRLVNPG GKDAVKDVQQKQ | 0.575 | 0.353 |
| AP02141 | LLRHVVKILEKYL | 1.531 | 0.682 |
| AP02159 | FFGHLYRGITSVVKHVHGLL SG | 1.144 | 0.4 |
| AP02163 | GFFGNTWKKIKGKADKIMLK KAVKIMVKKEGISKEEAQAK VDAMSKKQIRLYLLKYYGKK ALQKASEKL | 0.258 | 0.233 |
| AP02166 | LVPLFLSKLICFITKKC | 0.602 | 0.439 |
| AP02172 | FFGSLLSLGSKLLPSVFKLF QRKKE | 0.958 | 0.406 |
| AP02174 | FLPFLIPALTSLISSL | 0.898 | 0.331 |
| AP02179 | LNWGAILKHIIK | 0.881 | 0.585 |
| AP02197 | PAAAAQAVAGLAPVAAEQ | 0.425 | 0.31 |
| AP02202 | RKCNFLCKLKEKLRTVITSH IDKVLRPQG | 0.851 | 0.427 |
| AP02213 | GILGKLWEGFKSIV | 1.110 | 0.483 |
| AP02214 | IFGAIWKGISSLL | 1.066 | 0.435 |
| AP02215 | FLSTIWNGIKSLL | 1.239 | 0.487 |
| AP02216 | FLGALWNVAKSVF | 1.024 | 0.445 |
| AP02217 | FLSTLWNAAKSIF | 1.132 | 0.454 |
| AP02218 | IFKAIWSGIKSLF | 1.224 | 0.489 |
| AP02222 | FIPLVSGLFSRLL | 1.014 | 0.572 |
| AP02241 | KYALMKKIAELIPNLKSRQV K | 0.679 | 0.477 |
| AP02249 | GLLSLLSLLGKLL | 0.982 | 0.443 |
| AP02263 | YPELQQDLIARLL | 0.746 | 0.638 |
| AP02264 | FLSGILKLAFKIPSVLCAVL KNC | 0.958 | 0.36 |
| AP02266 | GFWDSVKEGLKNAAVTILNK IKCKISECPPA | 0.656 | 0.324 |
| AP02268 | GLLDSVKEGLKKVAGQLLDT LKCKISGCTPA | 0.831 | 0.324 |
| AP02273 | FITGLIGGLMKAL | 0.960 | 0.406 |
| AP02281 | GVLDTLKNVAIGVAKGAGTG VLKALLCQLDKSC | 0.370 | 0.304 |
| AP02283 | SLFGTFAKMALKGASKLIPH LLPSRQQ | 0.681 | 0.355 |
| AP02287 | GLKEVAHSAKKFAKGFISGL TGS | 0.972 | 0.348 |
| AP02293 | GFMSKVANFAKKFAKGGVNA IMNQK | 0.811 | 0.358 |
| AP02294 | FIGALLRPALKLLAGK | 0.767 | 0.514 |
| AP02298 | GAFGDLLKGVAKEAGMKLLN MAQCKLSGKC | 0.379 | 0.316 |
| AP02304 | FLAGLIGGLAKML | 0.889 | 0.387 |
| AP02317 | ITIPPIVKNTLKKFIKGAVS ALMS | 0.776 | 0.372 |
| AP02318 | IKIPSFFRNILKKVGKEAVS LIAGALKQS | 0.833 | 0.383 |
| AP02319 | GIFPIFAKLLGKVIKVASSL ISKGRTE | 0.885 | 0.383 |
| AP02335 | ILSYLWNGIKSIF | 1.242 | 0.484 |

**Table S2** – Loose data set

| **Sequence ID** | **Sequence** | **Score** | **Prediction** | **Class** |
| --- | --- | --- | --- | --- |
| D1 | ALFSLASKVVPSVFSMVTKK | 0.789 | AMP | Designed |
| D2 | VVFRVASKVFPAVYCTVSKK | 0.784 | AMP | Designed |
| D5 | FLFGLASKVFPAVYCKVTRK | 0.610 | AMP | Designed |
| D6 | LSAVGKIASKVVPSVIGAFK | 0.985 | AMP | Designed |
| D7 | PVIGKLASKVVPSVFSMIKR | 1.119 | AMP | Designed |
| D9 | GLMSLVKDIAKLAAKQGAKQ | 0.755 | AMP | Designed |
| D15 | SALGRVASKVFPAVYCSITK | 0.972 | AMP | Designed |
| D22 | LGALFRVASKVFPAVISMVK | 1.121 | AMP | Designed |
| D23 | ALGKLASKVFPAVYCTISRK | 0.782 | AMP | Designed |
| D24 | GFIGKLASKVVPSVYCKVTG | 0.687 | AMP | Designed |
| D25 | PVVFSVASKVVPSLISALKR | 1.018 | AMP | Designed |
| D28 | FLGVVFKLASKVFPAVFGKV | 0.766 | AMP | Designed |
| D29 | PAVFKIASKVVPSVYCKVSR | 0.954 | AMP | Designed |
| D30 | GALFGLASKVFPAVFGAFKK | 0.768 | AMP | Designed |
| D31 | SAVGKLASKVFPAVFSMVTK | 0.955 | AMP | Designed |
| D33 | VKDLAKFIAKTVAKQGGCYL | 0.750 | AMP | Designed |
| D34 | GVVGKLASKVVPSVFGSFTK | 0.802 | AMP | Designed |
| D35 | LPVVFRVASKVFPALISKLT | 0.968 | AMP | Designed |
| D36 | SAVGSVASKVVPSLISKVTK | 0.862 | AMP | Designed |
| D39 | MKSIAKFIAKTVAKQGAKQG | 0.533 | Non-AMP | Designed |
| D42 | LPAVFKLASKVVPSVFGLVK | 1.024 | AMP | Designed |
| D43 | SFVFKLASKVVPSVFSALTR | 1.072 | AMP | Designed |
| D44 | SVIGKIASKVVPSVYCAISK | 0.928 | AMP | Designed |
| D45 | PVVGRVASKVFPAVIGLVKK | 1.067 | AMP | Designed |
| D51 | FLFRVASKVFPALIGKFKKK | 0.841 | AMP | Designed |
| D55 | LSFVGRVASKVVPSLISMIK | 1.174 | AMP | Designed |
| D56 | SALGRLASKVVPAVIGKVTT | 0.876 | AMP | Designed |
| D57 | LGVVGSLASKVVPAVISKVK | 0.782 | AMP | Designed |
| D62 | LPAVFKLASKVFPAVYCKAS | 0.757 | AMP | Designed |
| D63 | LPVLFKLASKVFPAVFSSLK | 0.956 | AMP | Designed |
| D65 | VVGRVASKVVPSLIGLFTTK | 0.788 | AMP | Designed |
| D69 | SVVFGVASKVVPSVIGKVKT | 0.694 | AMP | Designed |
| D75 | FLPFVGRIASKVVPSVIGKV | 0.936 | AMP | Designed |
| D77 | GKKLAKTIAKEVAKQGAKFA | 0.890 | AMP | Designed |
| D82 | FVGSLASKVVPSVFGAIKTK | 0.664 | AMP | Designed |
| D83 | LPVVFKIASKVVPSVISKIT | 0.927 | AMP | Designed |
| D84 | GAVFGVASKVVPSVFSAIKK | 0.905 | AMP | Designed |
| D85 | FVGGVASKVVPSVYCKVSKK | 0.538 | Non-AMP | Designed |
| D88 | VVFKLASKVVPSVYCTITKK | 0.728 | AMP | Designed |
| D96 | GALFSLASKVVPAVIGLIKK | 0.895 | AMP | Designed |
| S1 | MVVFSVPKFKSTVAKLLSSA | 0.751 | AMP | Shuffled |
| S2 | TAKVVVFVSFSYVVPKKRAC | 0.205 | Non-AMP | Shuffled |
| S5 | FLPVLVKVFRYSKKTAAGCF | 0.838 | AMP | Shuffled |
| S6 | GVSSPIVAVKFKGAVASLIK | 0.290 | Non-AMP | Shuffled |
| S7 | SRVPLKSPVKIVGSKVMIFA | 0.368 | Non-AMP | Shuffled |
| S9 | GLKKDALQSIVKKAQLAAMG | 0.513 | Non-AMP | Shuffled |
| S15 | LYSPTCVKAAVSRFIGKVSA | 0.564 | Non-AMP | Shuffled |
| S22 | SVPSVGAVLFFKRAAVMKLI | 0.230 | Non-AMP | Shuffled |
| S23 | KYGPALVIAVKKSCSLTFRA | 0.650 | AMP | Shuffled |
| S25 | KSPFVLVVSSRVAAVIKSLP | 0.451 | Non-AMP | Shuffled |
| S28 | GVSVAGAKKVKVLFVFPFLF | 0.165 | Non-AMP | Shuffled |
| S29 | KVYVVKIAVPCFPKSARSVS | 0.684 | AMP | Shuffled |
| S31 | FMKVLAVFGSVVTSAPKASK | 0.790 | AMP | Shuffled |
| S33 | ALVYAGIKKTAFLKVQKCDG | 0.187 | Non-AMP | Shuffled |
| S34 | SVKPVGSSVVKGTALVKFFG | 0.517 | Non-AMP | Shuffled |
| S35 | KVFIATLVVSSFLLAKPPRV | 0.348 | Non-AMP | Shuffled |
| S36 | STVKVASKLAVVVSPISKGS | 0.611 | AMP | Shuffled |
| S39 | AKKAQKSGAQTIVKIFAKGM | 0.753 | AMP | Shuffled |
| S42 | VVAKKFFVLVKGLAPVLSPS | 0.667 | AMP | Shuffled |
| S43 | ASPTVFRSSVFLSLFVVAKK | 0.036 | Non-AMP | Shuffled |
| S44 | IASAVPVCVKGKISKSYISV | 0.201 | Non-AMP | Shuffled |
| S45 | VKRAGKGVAVVPSPLFKIVV | 0.968 | AMP | Shuffled |
| S51 | RKVAPALIKSFVFLFKFKKG | 0.728 | AMP | Shuffled |
| S55 | SSSIPIKMVLVRALVFVKSG | 0.369 | Non-AMP | Shuffled |
| S56 | TLVGVVAKLVATKIGSSPRA | 0.699 | AMP | Shuffled |
| S57 | PKVVGLSIVVVKAKVSSALG | 0.300 | Non-AMP | Shuffled |
| S62 | PSLLYKAKAVFCKPSAVAVF | 0.286 | Non-AMP | Shuffled |
| S63 | VSVKKVLPFAPLKSLLSFAF | 0.071 | Non-AMP | Shuffled |
| S65 | FKVVISKPGLSVRVGTALVT | 0.558 | Non-AMP | Shuffled |
| S69 | VFSVKGGKPSVVIKVVVAST | 0.151 | Non-AMP | Shuffled |
| S75 | SKFPLAGIFSVPGVKRVVVI | 0.370 | Non-AMP | Shuffled |
| S77 | VIAFAKTKEAKAKLKGQAKG | 0.133 | Non-AMP | Shuffled |
| S82 | KTVPVVLKASIKVSSAGFGF | 0.563 | Non-AMP | Shuffled |
| S83 | KIVKVITVKSISPASLVPVF | 0.393 | Non-AMP | Shuffled |
| S84 | SVKVAKSVIPSAVFAGGKVF | 0.674 | AMP | Shuffled |
| S85 | KVGKGSYPCSFVKVVAKVSV | 0.138 | Non-AMP | Shuffled |
| S88 | VKTKCSVPAVVYILVKTFKS | 0.533 | Non-AMP | Shuffled |
| S96 | LPVLFSSAIAKVGIKLGAKV | 0.387 | Non-AMP | Shuffled |

**Table S3** – Nagarajan data set

| **Sequence ID** | **Sequence** | **Score** | **Prediction** | **Class** |
| --- | --- | --- | --- | --- |
| NN2_0000 | EVAKKLLASALKLALAI | 0.877 | AMP | AMP |
| NN2_0001 | EDWNHLGAAVHTLKHVYK | 0.743 | AMP | AMP |
| NN2_0002 | AIVEQLRKRC | 0.898 | AMP | AMP |
| NN2_0003 | KLSASLKHVAHRARHLS | 0.935 | AMP | AMP |
| NN2_0004 | ESRAGKLAAKAAFKAAKR | 0.889 | AMP | AMP |
| NN2_0005 | EWAAARQVIIHATRKY | 0.544 | Non-AMP | AMP |
| NN2_0006 | EILSKALSALSPLAN | 1.150 | AMP | Non-AMP |
| NN2_0007 | EKAILSALKLLRLAL | 0.939 | AMP | AMP |
| NN2_0008 | ETAKGVAKHLPPAIA | 0.937 | AMP | Non-AMP |
| NN2_0009 | KVYARLHAVIKRLHRRLH | 1.332 | AMP | AMP |
| NN2_0018 | YLARAIRRTLARLLL | 1.254 | AMP | AMP |
| NN2_0022 | EWRVARRAVQRLRHLARRYH | 1.405 | AMP | AMP |
| NN2_0024 | ALKKMLRLAKRLS | 1.559 | AMP | AMP |
| NN2_0027 | VLSAFHKVIKIIHHISHF | 1.156 | AMP | AMP |
| NN2_0029 | RKFRKILHRARKWI | 1.707 | AMP | AMP |
| NN2_0035 | RRWGRWHRMRRRGR | 0.654 | AMP | AMP |
| NN2_0039 | FWKGLVKAAFKIVHAGS | 0.954 | AMP | AMP |
| NN2_0046 | GWKAIHKAAKGIHTYVN | 1.174 | AMP | AMP |
| NN2_0050 | SWKKFFKKARSLPKLF | 1.118 | AMP | AMP |
| NN2_0055 | YKRWKKWRSKAKKIL | 0.830 | AMP | AMP |
